# scFAIR Consortium: a decentralized hub for single-cell RNA-Seq data standardization and unification

**DOI:** 10.64898/2026.06.05.730084

**Authors:** Vincent Gardeux, Sara Carsanaro, Wen J. Chen, Fabrice P. A. David, Damien Goutte-Gattat, Jason A. Hilton, Tiago Lubiana, Nikhil Patel, Brian Raymor, Ida Zucchi, Bart Deplancke, Christina Ernst, David Osumi-Sutherland, Marc Robinson-Rechavi, Paul W. Sternberg, Frederic B. Bastian

## Abstract

The rapid accumulation of single-cell RNA-Seq (scRNA-seq) data across multiple repositories presents major challenges for data accessibility, integration, and reproducibility. While primary repositories provide raw data, they rarely include structured cell-type annotations or descriptions of analytical workflows, limiting the ability to reuse and integrate datasets in a FAIR (Findable, Accessible, Interoperable, Reusable) manner. Here we present scFAIR, a consortium of single-cell data resources that has developed a unified metadata schema and common curation framework to improve the FAIRness of scRNA-seq data. Building on and extending the CZ CELLxGENE Discover metadata schema, the scFAIR consortium has been instrumental in driving key schema improvements, including the expansion of supported organisms, richer biological context, and structured reporting of computational workflows. To provide unified access to decentralized datasets, the consortium developed the sc-fair.org portal, which currently aggregates 2,346 datasets across partner resources through ontology-aware semantic search. We demonstrate the practical value of FAIR-compliant datasets through a cross-species validation between human and mouse Allen Brain Atlases, showing that standardized ontology annotations enable reliable annotation transfer across species, with 90% of neuronal clusters receiving an exact or equivalent label. Together, the scFAIR schema, validator, and portal constitute a community-driven framework that advances single-cell data standardization and lays the foundation for reproducible, large-scale integration of single-cell datasets.

## Introduction

In recent years, single-cell RNA sequencing (scRNA-seq) has revolutionized our understanding of cellular heterogeneity, providing unprecedented insights into the functional diversity of individual cells within complex tissues (see, e.g., (1, 2)). However, the rapid accumulation of scRNA-Seq data across multiple repositories presents significant challenges in data accessibility, integration, and reproducibility. Ensuring that these data adhere to the FAIR (Findable, Accessible, Interoperable, Reusable) principles(3) is critical for advancing scientific discovery and enabling robust, reproducible research.

Currently, most scRNA-Seq datasets live in primary repositories such as GEO(4), ArrayExpress(5), or SRA(6), where raw data are provided, but, in most cases, not associated with structured information about cell types, and with no description of the analyses performed. This limits our ability to make these datasets reusable and interoperable. Some knowledgebases and platforms, such as CZ CELLxGENE Discover(7) (referred throughout as CxG forward), ASAP(8) or Bgee(9), provide structured information about, e.g., cell types identified by authors, using ontologies like the Cell Ontology(10). This makes the datasets highly queryable and interoperable. Other resources, including the Broad Institute Single Cell Portal(11) and the UCSC Cell Browser(12), aim at supporting researchers for exploring and visualizing scRNA-Seq datasets, but do not aim at data standardization. As a result of these different efforts, researchers have multiple options to make their data accessible, ranging from basic access to raw datasets, to interactive visualizations, and ultimately to highly structured, curated information resources. Researchers are thus faced with a fragmented ecosystem in which datasets are dispersed across multiple platforms, each offering different levels of structure, curation, and accessibility, with no straightforward way to identify related datasets, to integrate them across repositories, or to make them interoperable.

Recognizing these challenges, recent efforts have emphasized the need for standardized metadata describing not only biological annotations but also the analytical processes that generate single-cell data. Notably, the Matrix and Analysis Metadata Standards (MAMS) initiative(13) proposes a formal schema for describing expression matrices and their associated analytical workflows, including matrix types, annotation classes, and analysis provenance such as software tools and parameters. By explicitly capturing how data matrices and derived annotations are produced, MAMS addresses a critical gap in current practices, where analytical provenance is often inconsistently reported or lost when datasets are shared or reanalyzed. This work highlights the importance of workflow-aware metadata as a prerequisite for reproducibility and large-scale integration, and provides a complementary framework to repository-focused standardization efforts.

In parallel, recent discussions of best practices for atlas construction have been synthesized(14), outlining principles for dataset diversity, batch correction, annotation, ontology alignment, evaluation, and FAIR accessibility. This work represents a reference framework for large-scale single-cell integration projects. Importantly, the authors emphasize that ontological consistency, transparent annotation, and reproducible metadata are prerequisites for making atlases reusable across studies.

The scFAIR initiative directly builds on these considerations by not only adopting common metadata schemas and ontologies, but also validating this framework through a cross-atlas validation as a systematic quality-control step. Therefore, scFAIR translates the conceptual guidelines of Hrovatin et al.(14) into practical workflows and tools that ensure annotation transferability and reproducibility across resources.

scFAIR is a consortium of resources providing added-value curation of scRNA-Seq data, and was funded by members of Alliance of Genome Resources (AGR)(15), ASAP(8), Bgee(9), CxG(7), EMBL-EBI Single Cell Expression Atlas(SCEA)(16), FlyBase(17), and contributors to the Cell Ontology (CL)(18), with the aim of improving the FAIRness of this type of data. scFAIR is onboarding more resources and connecting to other initiatives, such as the ELIXIR Single Cell Omics Community(19). The consortium agreed on a common metadata schema for curation, defined adaptors to individual resources for improving data retrieval, and tackled the problem of analysis reproducibility. We present here the result of this work and the recommendations to streamline the process of data sharing and analysis, ultimately fostering a more collaborative and efficient scientific environment. To support this effort, we also created the sc-fair.org portal, a website that centralizes our findings, and aggregates decentralized datasets based on our common metadata schema.

### Current state of single-cell metadata standardization efforts

The standardisation of single-cell metadata is a continuation of decades of community effort in biology, and more specifically in high-throughput sequencing data sharing. It draws directly from the foundations of MIAME (Minimum Information About a Microarray Experiment)(20) and MINSEQE (Minimum Information About a High-Throughput Nucleotide Sequencing Experiment)(21), which established the requirements for archival integrity in microarray and bulk RNA-Seq sequencing. As the field transitioned to single-cell resolutions, the Human Cell Atlas (HCA) initiative(22) performed pioneering work on unifying metadata standards through a transparent, community-driven approach. In collaboration with EMBL-EBI, the minSCe (Minimum Information about a Single-Cell Experiment) guidelines(23) was developed and implemented within ArrayExpress. Building on the principles of MINSEQ, minSCe ensures the preservation of unique single-cell complexities, such as tissue dissociation protocols, cell enrichment methods, and barcode configurations. By capturing these attributes, the framework ensures that the full experimental lineage remains available for robust re-analysis and reproducibility.

While these archival standards ensure data provenance and analysis reproducibility, the rapid growth in secondary analysis and interactive visualisation necessitated a more “query-ready” framework. The CxG metadata schema emerged as a widely adopted implementation of these broader community principles. CxG is an interactive data explorer designed for visualizing and analyzing scRNA-seq data, developed by the Chan Zuckerberg (CZ) Initiative. As part of its visualization capacities, CxG hosts hundreds of public scRNA-seq datasets submitted by research labs to make their data broadly available to the community. As CxG relies on motivated contributors to submit datasets, there is pressure to keep submission burden to a minimum. This makes CxG datasets highly useful for exploratory analyses and cross-study comparisons, but potentially less comprehensive for reproducing the full experimental workflow.

The CxG schema spans multiple hierarchical levels: dataset, cell, and gene. At the dataset level, the schema defines identifiers and versioning to allow stable referencing and tracking of updates. It also captures descriptive information such as dataset titles, detailed summaries, and study descriptions, along with publication DOIs which provide links back to the primary research context, principal investigators, and institutional affiliations. At the cell level, annotations include barcodes and unique identifiers as well as biological attributes such as cell type labels, which are often harmonized against ontologies like CL(18) or UBERON(24), or taxon-specific ontologies e.g. FBbt(25) for *Drosophila*. The schema also accommodates experimental conditions including tissue of origin, developmental stage, and disease status, all of them being encoded by different ontologies. For example, disease conditions are encoded via the Mondo Disease Ontology (MONDO), enabling fine-grained and interoperable clinical annotations. Beyond scRNA-seq, the schema is designed to accommodate multiple data modalities: it defines specific matrix layer requirements for scATAC-seq (both unpaired and paired with RNA), as well as for spatial transcriptomics assays such as Visium and Slide-seqV2, and includes support for perturbation-based assays such as sci-Plex. In addition, other types of information can also be recorded, even if not mandatory at the moment, such as sequencing platform, library preparation, or quality-control related metrics. At the gene level, the schema specifies identifiers such as Ensembl IDs and gene symbols, together with functional annotations including canonical names, aliases, and cross-references to standardized resources like FlyBase.

These metadata elements are tightly integrated with the expression matrix, which remains the core component of every dataset. The matrix stores gene expression values for individual cells and is most commonly provided in the H5AD format. Indeed, H5AD has emerged as the dominant choice over alternatives such as Loom or the 10x H5 format, primarily because its structured design allows the storage of multiple layers of metadata alongside the raw expression data within a single file.

To support adoption of the metadata schema, CxG provides a dedicated validation tool that programmatically checks dataset submissions for compliance with required fields, data types, and ontology constraints. This validator has become an important component of the submission workflow where it provides immediate feedback to data providers prior to publication. By enforcing schema consistency at submission time, the validator plays a critical role in improving baseline data quality and interoperability across resources. However, the validator’s scope is intentionally limited to syntactic and structural correctness. While it can verify that required fields are present, values conform to expected formats, and ontology terms are valid, it cannot assess semantic correctness, biological plausibility, or the adequacy of experimental and analytical descriptions. For example, the validator cannot determine whether a chosen cell type annotation is appropriate, whether preprocessing steps are sufficiently documented, or whether important sources of biological variation are missing. As such, automated validation must be complemented by additional layers of quality assurance. While CxG provides expert curation focused on metadata accuracy within the scope of its minimal schema, community-agreed best practices are needed for preprocessing documentation and experimental design reporting areas where scFAIR aims to provide guidance.

When evaluated against the FAIR principles, the strengths and gaps of the CxG schema become clear. Datasets are **Findable**, with stable identifiers and cross-references to publications; they are **Accessible**, hosted through an open portal with standardized formats; and they are **Interoperable**, thanks to ontology alignment and the consistent use of H5AD. However, **Reusability** remains incomplete, as insufficient reporting of experimental design, raw data processing, and multi-modal integration limits the ability to fully reproduce studies. Moreover, the CxG schema remains limited in the accepted species (constrained by the data types formally supported by CxG), genome and genome annotation versions used (pinned directly in the schema at any given version), or accepted ontology terms (with specific ontologies assigned for each accepted species).

Bridging this gap thus requires extending CxG metadata toward richer, workflow-aware descriptions. We present here the scFAIR extension of the CxG metadata schema, with its accompanying updated validator.

### scFAIR metadata schema

The scFAIR metadata schema builds directly on the CZI CxG schema, while adapting it to the needs of a broader, federated single-cell ecosystem (**Fig 1**). Rather than defining an entirely separate standard, scFAIR preserves compatibility with the widely adopted H5AD/AnnData-based representation used by CxG, while extending the schema in areas that are essential for cross-resource discovery, multi-species reuse, and reproducibility (**Table 1**). The current scFAIR schema is versioned and is organized into a modular set of documents: a core schema covering shared single-cell metadata, together with dedicated extensions for analysis provenance, spatial assays, perturbation experiments, and scATAC-seq data. This modular organization allows scFAIR to maintain a stable common metadata layer while enabling modality-specific requirements to evolve independently.

**Table 1.** Fields updated in scFAIR metadata schema.

| Field | CxG schema v 7.1.0 requirement | scFAIR schema v 7.1.0+scFAIR1.0 |
| --- | --- | --- |
| obs H5AD layer (Cell metadata) |  |  |
| development_stage_ontology_term_id | Human: HsapDv<br>Mouse: MmusDv<br>Fly: FBdv<br>Zebrafish: ZFS<br>C. elegans: WBIs<br>Other organisms: any term descendant of<br>UBERON:0000105 « life cycle stage » | Terms from any ontology part of of the developmental stage ontology repository (27) |
| tissue_ontology_term_id | Fly: FBbt or Uberon<br>Zebrafish: ZFA or Uberon<br>C. elegans: WBbt or Uberon<br>Other organisms: Uberon | Any term part of the composite or collected metazoan version of Uberon |
| strain_or_genetic_background_term_id and strain_or_genetic_background | NA | Additional column to capture EFO or free text information about strain, stock, line, cultivar, genotype, ecotype, variety, serovar, or breeding and crossing design for any non-human organisms |
| var and raw.var H5AD layer (Gene Metadata) |  |  |
| gene_id | specific assembly and gene annotation version depending on the schema version, in 14 species + SARS-CoV-2, as of schema v. 7.1 | Schema should allow users to specify the assembly and gene annotation version used |
| uns H5AD layer (Dataset Metadata) |  |  |
| analysis_pipeline | NA | A schema-constrained JSON entry describing the analysis pipeline used to preprocess and perform the downstream analysis of the dataset. |
| ensembl_release | NA | Ensembl release number of the assembly used for gene annotation |
| ensembl_database | NA | Ensembl database name such as EnsemblMetazoa |
| ensembl_assembly | NA | Ensembl assembly name of the assembly used for gene annotation, e.g. "GRCh38.p14" for Homo Sapiens release 115 |

**Figure 1.**
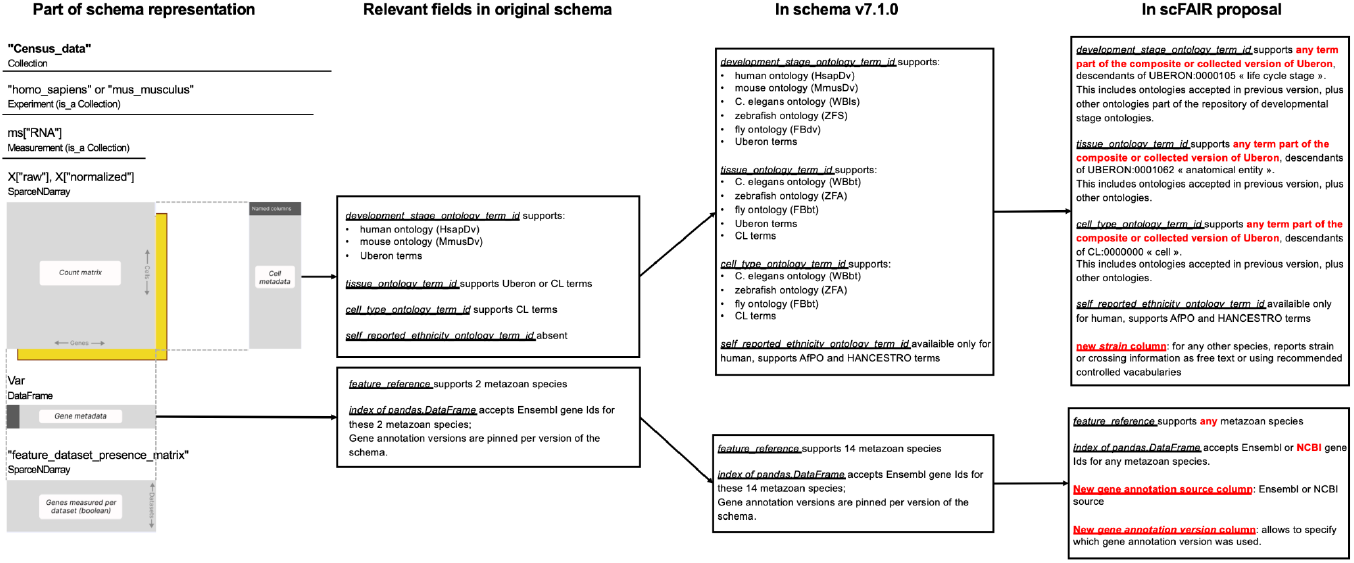
part of CxG metadata schema representation, highlighting fields relevant to the scFAIR update. Red: scFAIR update. Adapted from https://chanzuckerberg.github.io/cellxgene-census/_images/census-spatial-schema.svg with permission.

A central contribution of scFAIR is the removal of species-specific bottlenecks that limited earlier single-cell metadata schemas. Whereas previous CxG schema versions were tied to a defined set of supported organisms and reference annotations, scFAIR generalizes the representation of organism and gene metadata by allowing any species represented in the Ensembl and Ensembl Metazoa ecosystem. To support this broader taxonomic scope, the schema introduces explicit dataset-level fields for the Ensembl database, release, and assembly used for gene annotation. This is important because the biological interpretation of an expression matrix depends not only on the gene identifiers present in the final object, but also on the genome assembly and annotation release used during quantification. By making this information explicit, scFAIR reduces ambiguity in downstream reanalysis and supports more reliable comparison of datasets generated across species, assemblies, and annotation versions.

scFAIR also extends ontology-based annotation beyond the constraints of a small number of taxon-specific configurations. For anatomical entities, cell types, and life and developmental stages, the schema recommends use of the collected or composite metazoan versions of Uberon(24). The collected ontology merges Uberon itself with the CL ontology, and several taxon-specific ontologies (e.g., FBbt(25), ZFA(26), FBdv, HsapDv(27)), describing anatomy, and developmental and life stages. The composite ontology is derived from the collected ontology, by removing, wherever possible, the taxon-specific terms, and mapping them to the corresponding taxon-neutral terms from Uberon, resulting in less redundancy (see Combined Multispecies Ontologies document for more information). This approach preserves the advantages of taxon-neutral terms where they are sufficiently precise, while allowing species-specific terms when they provide the most accurate biological description. In practice, this makes the schema better suited to comparative biology: datasets can be queried through shared ontology structure, while still retaining biologically meaningful species-specific annotations where needed. Of note, the related taxon-specific ontologies can still be directly used, as they are referenced in the collected- and composite-metazoan versions of Uberon.

For human datasets, the schema preserves the existing *self_reported_ethnicity_ontology_term_id* field, which may be populated using AfPO or HANCESTRO terms, or reported as “unknown” or “na” where appropriate. This field follows the current CxG convention and is treated as a human-specific descriptor. scFAIR does not extend this field to non-human organisms and does not reinterpret it as a general measure of genetic background.

Separately, for non-human organisms, scFAIR introduces a dedicated *obs* level field, *strain_or_genetic_background*, to record experimentally relevant biological background information when available. This field may capture strain, stock, line, cultivar, genotype, ecotype, variety, serovar, or breeding and crossing design, depending on the organism and community practice. A companion field, *strain_or_genetic_background_ontology_term_id*, may be used when an appropriate controlled term exists, with EFO recommended as a general cross species source and organism specific nomenclatures or ontologies used where available. When no suitable ontology term exists, the free text field should still be populated in a standardized manner. This metadata is modeled independently from human specific descriptors and is intended to improve reproducibility by making experimentally relevant genetic background explicit.

These descriptors are important for reproducibility because genetic background can influence baseline expression, phenotype, and response to perturbation in model organisms. Indeed, in many model systems, particularly mouse, zebrafish, or rat, strain background is a major source of transcriptional variation and experimental reproducibility. For example, well-documented differences between commonly used strains such as C57BL/6J and BALB/c affect immune responses, metabolic phenotypes, and baseline gene expression profiles(28, 29). In addition, and similarly to recent versions of the CxG schema, to which the scFAIR consortium directly contributed, scFAIR allows support for cell lines defined in the Cellosaurus database (30), enabling unambiguous identification of in vitro models across studies. By linking cell line annotations to this widely adopted reference, the schema now allows transparent tracking of cell line provenance, which is particularly valuable for perturbation screens and disease modeling.

A second major extension introduced by scFAIR concerns analysis provenance. The core schema adds an optional *analysis_pipeline* entry in *uns*, storing a JSON description of the computational workflow used to generate the dataset. The companion *schema_analysis_json*.*md* document defines how analysis steps should be described in execution order, including the step type, method or tool, command line or function call, software version, programming language, parameters, inputs, outputs, and computational environment. This provides a structured mechanism to report steps such as alignment, counting, quality control, normalization, dimensionality reduction, batch correction, integration, clustering, marker-gene detection, and cell type annotation. The schema does not require every dataset to be fully executable end-to-end, but it creates a shared framework for distinguishing undocumented analyses from analyses whose tools, parameters, and environments are sufficiently described to support reproducibility.

This workflow-aware extension is particularly important for single-cell data because small differences in preprocessing and analysis can substantially affect downstream results. Indeed, even when gene identifiers in the final expression matrix appear standardized, reproducing or reinterpreting a dataset can become difficult or impossible if genome assemblies, reference annotation versions, and processing steps are not explicitly documented. Key analytical choices, including the aligner or pseudoaligner, genome reference, filtering thresholds, normalization method, integration strategy, clustering resolution, and annotation procedure, can substantially affect the resulting cell populations, marker genes, and biological conclusions(31–33). By capturing these decisions in a machine-readable JSON structure, scFAIR provides a practical bridge between lightweight metadata submission and full computational reproducibility. At a basic level, the schema enables users and curators to understand how an object was produced; at a higher level, links to Docker images, image digests, or conda environments can support rerunning or auditing key workflow steps.

Together, these developments make the scFAIR schema a federated metadata standard for reusable single-cell data. It retains the practical strengths of the CxG model: H5AD compatibility, ontology-backed metadata, and validator-driven consistency, while adding the metadata needed for broader species coverage, reproducible analysis, and cross-resource integration. As partner resources adopt the schema and expose their datasets through sc-fair.org, these extensions provide the basis for a more coherent single-cell data ecosystem in which datasets can be discovered, compared, reanalyzed, and interpreted across repositories without sacrificing provenance or biological specificity.

### Unified view of decentralized datasets

The sc-fair.org portal is the central platform developed by the scFAIR consortium to provide a unified entry point to decentralized single cell RNA seq resources (**Fig 2**). Rather than duplicating existing repositories, the portal acts as an integration layer that aggregates datasets, metadata, and ontology information across partner platforms, while preserving the original hosting environments and attribution to data providers (**Fig 2A**). This federated design reflects the core philosophy of scFAIR: improving findability, interoperability, and reuse without centralizing data ownership.

**Figure 2.**
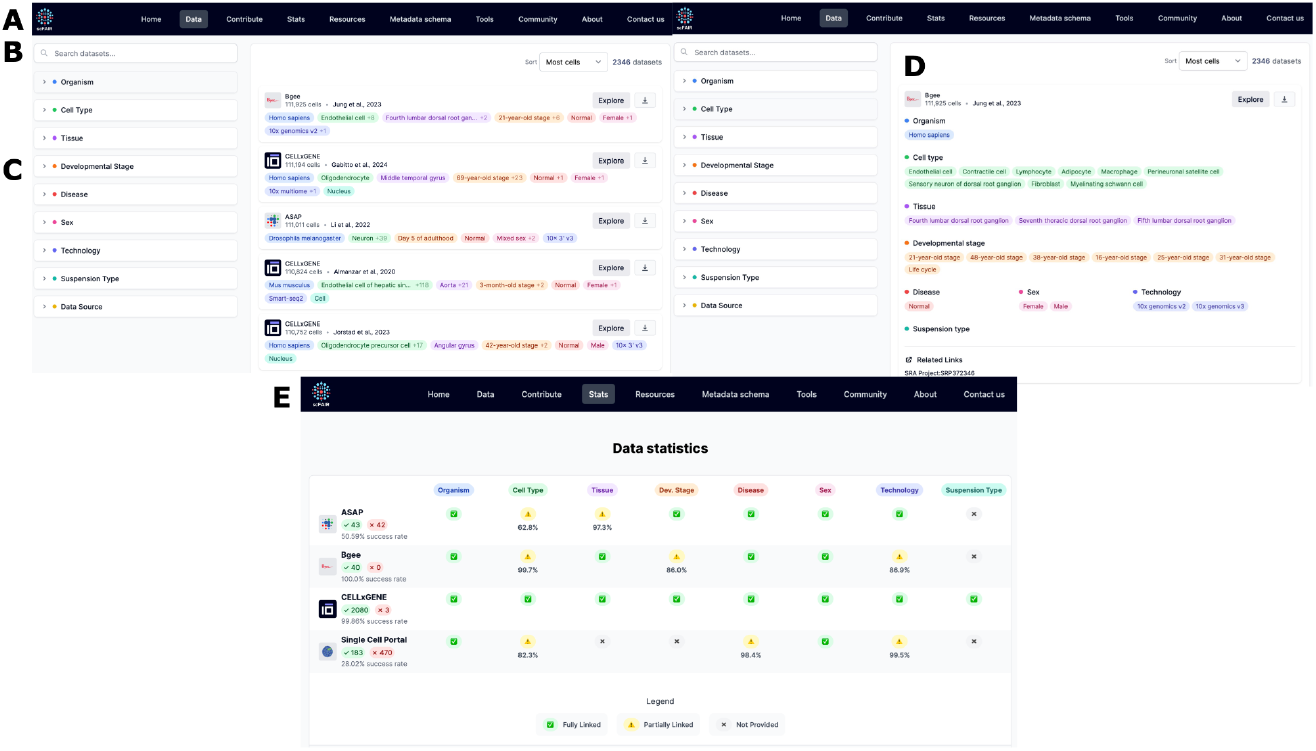
Single-cell data aggregation and FAIR compliance. **A.** the data tab shows the aggregate of scRNA-Seq datasets across scFAIR consortium partners; **B**. the search function leverages the hierarchical ontologies to return all relevant results; **C**. the filter function allows datasets to be filtered by various high-impact criteria; **D**. expanding a dataset displays an overview of annotations present in the dataset; **E**. the stats tab shows datasets and data types that are scFAIR compliant, stratified by scFAIR partner resource.

CxG, Bgee, and ASAP were the first partner resources programmatically integrated into the portal. For each resource, dedicated API adapters retrieve and normalize metadata at regular intervals, allowing the portal to remain synchronized with the source databases while minimizing manual curation effort. Dataset integrity and update tracking are supported through SHA 256 hashing, which enables change detection and helps prevent data drift across synchronization cycles. This architecture allows the portal to scale with the rapid growth of public single cell datasets while preserving provenance from each contributing resource.

A key feature of the portal is its ontology aware search system (**Fig 2B**). The platform parses ontology structures and builds adjacency tables that capture parent and child relationships for terms describing tissues, anatomical structures, developmental stages, and cell types (**Fig 2C-D**). As a result, users can retrieve datasets not only by exact annotation terms, but also through broader or more specific ontology relationships. For example, a query for “brain” can recover datasets annotated to more specific brain regions. This functionality turns the portal from a simple dataset index into a semantic search engine for FAIR single cell data (**Fig 2B-D**).

The portal also includes a statistics page (**Fig 2E**) that provides an overview of metadata completeness and schema compliance across integrated resources. For each partner resource, the page reports the number of datasets that pass validation against the latest schema, together with success rates and field level summaries for key metadata categories such as organism, cell type, tissue, developmental stage, disease, sex, technology, and suspension type. It also distinguishes metadata values that are fully linked to ontology terms, partially linked because of parsing issues, or not provided. In the current portal, validation rates vary substantially across resources, reflecting differences in metadata models, curation practices, and available API fields. For example, CxG datasets show near complete validation against the schema, whereas other resources expose complementary strengths, broader species coverage, or different levels of ontology linkage. By making these differences visible, the stats page turns schema compliance into a measurable and transparent property of each resource, helping identify where harmonization is already effective and where additional curation, mapping, or schema alignment is needed.

The portal also provides a practical framework for cross resource metadata harmonization. Because each partner resource exposes metadata through its own API and internal data model, scFAIR currently relies on resource specific adapters to map external metadata into the shared scFAIR representation. This approach enabled rapid onboarding of initial partners, but it also highlights the need for a common API schema that future resources could implement directly. Ongoing work to map the SCEA metadata schema to the scFAIR schema represents an important step in this direction. In the longer term, a shared API specification would reduce the need for custom adapters, support richer queries across resources, and facilitate the assignment of stable scFAIR level accessions while preserving provenance across platforms.

The consortium is continuing to expand the portal through additional integrations, including SCEA and the Broad Institute Single Cell Portal. Beyond dataset aggregation, the portal also functions as a community hub: it hosts the agreed scFAIR metadata schema, provides guidance for dataset curation, and highlights ongoing FAIR initiatives across the single cell ecosystem. In doing so, it connects technical infrastructure with community practice, supporting researchers in both locating and responsibly reusing single cell data.

A current limitation of the portal is the presence of duplicated or partially overlapping datasets across integrated resources. The same primary study may be represented in multiple portals, but these entries are not always identical, because they may reflect different preprocessing, filtering, reannotation, or submission versions. Merging such records automatically could obscure meaningful differences between derived datasets. For this reason, the portal currently preserves resource specific entries and provenance, while future work will aim to represent dataset relationships more explicitly, distinguishing true duplicates from distinct reanalyses or versions of the same underlying experiment.

### Validation using the Allen Brain Atlases

To evaluate the interoperability of scFAIR-compliant resources, we performed a cross-atlas validation between the human Allen Brain Atlas (34) and our curated mouse Allen Brain Atlas(35). Both datasets adhere to FAIR principles, particularly in their use of standardized ontologies and structured file formats, enabling annotation transfer through SAMap(36). This experiment tested whether annotations could be reliably propagated between species and, critically, whether FAIR-compliant datasets ameliorate the process of data integration and analysis, and assist in identifying biologically relevant findings.

We first applied SAMap to align the 382 neuronal human clusters against the mouse brain atlas (**Fig 3A,B**). We next removed clusters with imprecise annotations where we did not expect a match, leaving 270 clusters for the analysis. We found that 56% of the clusters were labeled with the exact ontology term that was originally applied. The use of a unified multi-species ontology allowed for the effective transfer and validation of cell labels, despite transferring from one species to another. For instance, inhibitory neuron subtypes, astrocytes, and microglia showed robust one-to-one mappings between human and mouse datasets, with alignment scores exceeding 0.95 in most cases.

**Figure 3.**
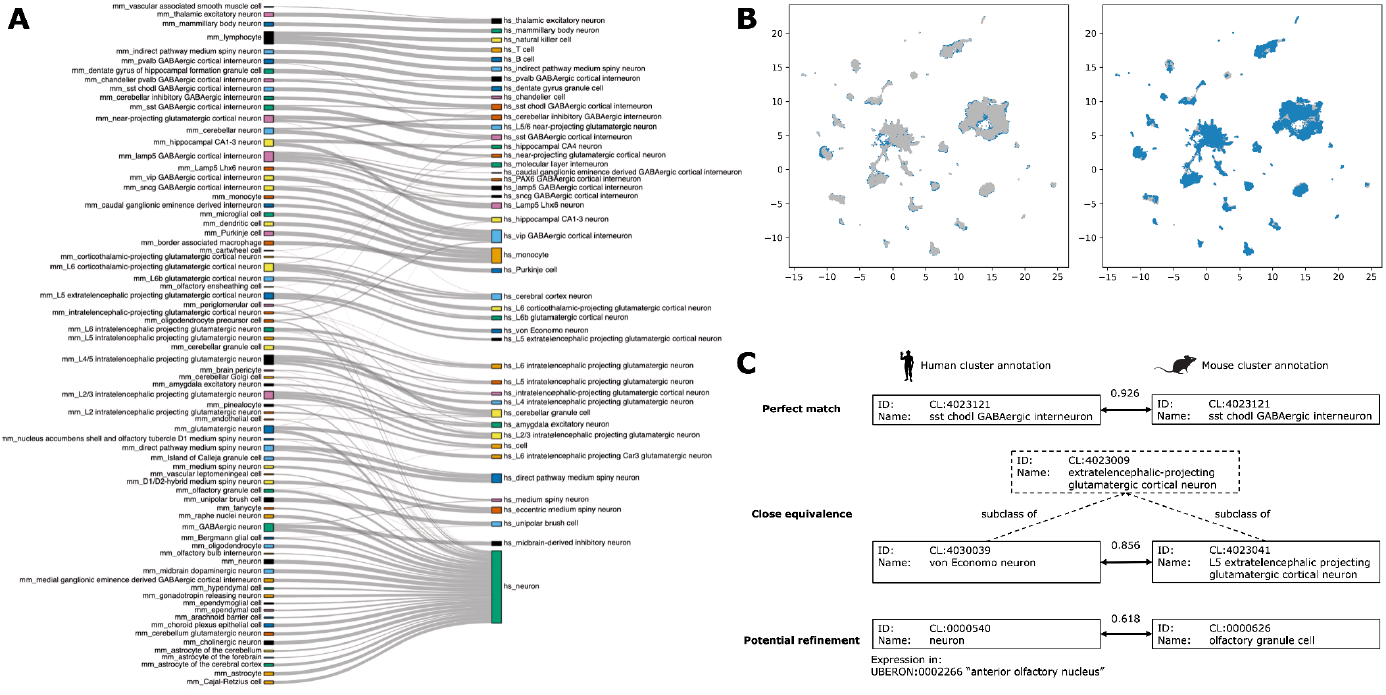
Results of SAMap mapping between mouse and human Allen Brain Atlas. **A.** Sankey plot of the aligned clusters between mouse (left) and human (right). **B**. UMAP projection of the combined human and mouse clusters; grey: human cells; blue: mouse cells; left panel: human cells on top; right panel: mouse cells on top. **C**. Examples of annotation perfect match, of close equivalence, and of mismatch where additional information could be used to refine the annotation Solid line boxes: cluster annotations in human (left) and mouse (right); dash lines: additional superclass information in the Cell Ontology; double arrows: SAMap alignment scores.

Next, we reviewed the mismatched clusters and used ontology reasoning, plus manual checks when the ontology structure did not satisfy the proper composition of terms, to identify equivalent terms. We found that 90% of clusters were labeled with either an exact or equivalent label, encompassing 150 exact matches and 92 equivalent matches (**Fig 3C**). This high concordance demonstrates the feasibility of transferring annotations across FAIR-compliant atlases.

The ontology structure allowed us to efficiently identify equivalent matches from intersecting parent-child, sibling, and species-specific terms, even for not conspicuously related terms (**Fig 3C**). The majority of them reflected differences in granularity or anatomical scope. For example, the mouse cluster annotated to CL:4030062 “L4/5 intratelencephalic projecting glutamatergic neuron” was applied to 3 human clusters labeled as CL:4023008 “intratelencephalic-projecting glutamatergic cortical neuron”: from the Cell Ontology, L4/5 neurons are equivalent to CL:4023040 “L2/3-6 intratelencephalic projecting glutamatergic neuron” (a subclass of CL:4023008 used in the human label), but more precisely located in “cortical layer IV/V” (UBERON:8440001); thus it was possible to automatically infer the close relation of the human and mouse terms. In another case, a cluster in mouse annotated to CL:4023041 “L5 extratelencephalic projecting glutamatergic cortical neuron” was mapped to the human cluster labeled CL:4030039 “von Economo neuron”; while those labels are not conspicuously related, both terms are siblings, subclasses of the term CL:4023009 “extratelencephalic-projecting glutamatergic cortical neuron”, thus the equivalence could also be inferred automatically. SAMap scores in these equivalent matches were generally more moderate, averaging 0.52, consistent with partial rather than exact match, due to the differences in granularity of cell identification, or brain location.

Finally, our analysis highlighted several clusters where the original annotation could be reassessed and possibly updated **(Fig 3C)**. For example, a human cluster annotated as “neuron” (CL:0000540), with high expression in the “anterior olfactory nucleus” (UBERON:0002266) was assigned the mouse label “olfactory granule cell” (CL:0000626) with an alignment score of 0.91. We also identified a human cluster annotated with the term cerebellar granule cell (CL:0001031) that was assigned the mouse label cerebellar Golgi cell (CL:0000119, alignment score 0.61). The cluster shows expression of inhibitory neurotransmitters, which supports the annotation of cerebellar Golgi cell, an inhibitory neuron, as opposed to cerebellar granule cells that function as excitatory neurons.

Together, these results confirm that FAIR-aligned resources such as the human Allen Brain Atlas and the mouse Allen Brain Atlas can be directly integrated through SAMap, yielding biologically meaningful annotation transfers. Beyond validating interoperability, the analysis highlights the added value of using standardized ontologies, and revealed specific opportunities for refining curation, including harmonizing granularity in anatomical definitions and critically reassessing ambiguous cluster assignments. Incorporating such cross-atlas validations into the scFAIR workflow thus serves both as a test of reusability and as a practical guide for ongoing annotation refinement. This approach would be even more advantageous for integrating more distant species thanks to the use of multispecies ontologies, without which equivalent terms are less obvious and the subsequent data analysis more subjective and labor-intensive.

## Discussion and future directions

### scFAIR contributions

The scFAIR consortium addresses three interconnected challenges that limit the reuse and integration of single cell RNA seq data: fragmented access across repositories, restricted species coverage in existing metadata schemas, and insufficient capture of analysis provenance.

First, by extending the CxG metadata schema, scFAIR broadens the range of organisms for which FAIR compliant single cell data can be produced and exchanged. The expansion from a primarily human and mouse centered schema toward support for a wider range of metazoan species, together with the adoption of composite Uberon resources for tissues, cell types, and developmental stages, removes an important barrier for comparative biology. The addition of explicit information about genome assemblies and gene annotation versions further improves interpretability and reproducibility. In this way, scFAIR provides a pragmatic and interoperable extension of CxG that improves findability, cross species comparison, and reuse while remaining compatible with existing community data flows.

Second, the sc fair.org portal demonstrates that a federated and ontology aware aggregation approach can surface datasets across decentralized repositories. By providing a unified semantic search interface over partner resources, without centralizing data ownership, the portal improves findability while preserving the original hosting environments and resource specific expertise. Hierarchical ontology querying enables researchers to retrieve datasets annotated at different levels of biological specificity, from broad anatomical structures or cell classes to fine grained tissues and cell types, supporting both discovery and comparative analyses.

Third, scFAIR lays the groundwork for improved analysis reproducibility through structured capture of computational workflows and parameters. The analysis metadata schema provides a common framework for recording key steps such as alignment, quality control, normalization, integration, clustering, and annotation, together with the tools, versions, parameters, inputs, outputs, and computational environments used. While full reproducibility remains an evolving community goal, this structured representation provides a concrete step toward making analytical decisions more transparent, comparable, and reusable across resources.

The cross species validation between the human and mouse Allen Brain Atlases illustrates the practical value of these developments. Using FAIR compliant, ontology standardized datasets, 90% of neuronal clusters received an exact or equivalent label through SAMap, and ontology structure enabled inference of equivalences even for terms that were not obviously related. This result demonstrates that standardized metadata and ontology based annotation can support reliable annotation transfer across species, and suggests that cross atlas validation could become a scalable quality control strategy for the single cell community.

### Challenges for analysis reproducibility

While recent advances in metadata standardization have improved the findability and interoperability of single cell RNA seq datasets, analysis reproducibility remains a major challenge. Current metadata schemas mainly capture biological annotations and high level experimental descriptors, but they often leave important computational details insufficiently documented. In practice, many datasets are generated through complex pipelines involving alignment or pseudoalignment, quality control, normalization, dimensionality reduction, integration, clustering, and annotation. Even when gene identifiers in the final expression matrix appear standardized, reproducing or reinterpreting a dataset can become difficult or impossible if genome assemblies, reference annotation versions, software versions, filtering thresholds, processing parameters, and annotation strategies are not explicitly recorded.

Small differences at any of these steps can substantially affect the resulting cell populations, marker genes, and biological conclusions(37). Yet these decisions are often described only narratively, if at all, and are rarely captured in a form that can be compared across tools or repositories. Some platforms already provide partial solutions. For example, the ASAP portal exports analysis workflows and parameters in JSON format. However, these representations have so far remained specific to individual platforms, limiting their reuse across the broader single cell ecosystem.

To address this gap, scFAIR introduces a structured JSON schema for analysis metadata. This schema provides a common representation for ordered analysis steps, including the type of step performed, the tool or method used, command line or function call, software version, parameters, input and output files, and computational environment. Existing resources such as the Experimental Factor Ontology (EFO)(38), together with registries of bioinformatics tools, could further support consistent naming of software, methods, and analysis concepts. At a basic level, this structured JSON description makes analytical decisions more transparent and comparable across datasets. At a higher level, links to container images, conda environments, or workflow managers such as Snakemake could support more complete rerunning of analyses.

Together, these efforts position reproducibility as an evolving, community driven goal rather than a single fixed requirement. scFAIR contributes to this goal by providing a practical framework for recording analysis provenance, while leaving room for resources and communities to progressively adopt richer levels of workflow documentation.

### Limitations in capturing additional sources of biological variation

Beyond technical and provenance metadata, current single cell schemas still capture only part of the biological variation that can influence gene expression and affect data reuse. Classical standards such as MIAME and MINSEQE emphasize the reporting of experimental design variables and biological conditions that may affect transcriptional outcomes. By contrast, current single cell metadata schemas generally encode a core set of well defined descriptors, such as developmental stage, biological sex, anatomical location, and disease status, while other relevant dimensions remain less systematically represented.

One important source of variation is genetic background in non-human organisms. For model organisms, information such as strain, stock, line, genotype, cultivar, ecotype, or crossing design can strongly influence baseline expression, phenotype, and response to perturbation. scFAIR therefore proposes to capture this information through a dedicated metadata field that is independent from human specific descriptors. Where possible, such information should be represented using established organism specific nomenclatures, controlled vocabularies, or ontologies such as EFO, complemented by standardized free text when no suitable term is available.

Another important limitation concerns the distinction between cell identity and cell state. Author provided annotations often describe transient or functional states, such as *proliferating type II pneumocyte* or *activated macrophage*, which do not always map cleanly onto the Cell Ontology representation of stable cell identities. These states could be captured more explicitly through controlled vocabularies such as the Gene Ontology, by recording biological processes that are active at the time of measurement and that drive systematic expression differences.

More broadly, additional contextual variables such as time of sampling, circadian effects, environmental and housing conditions, diet, medication exposure, treatment history, or post mortem interval are rarely recorded in a structured way, despite their influence on gene expression. These factors could in principle be formalized using phenotype and experimental ontologies such as the Experimental Factor Ontology and the Ontology for Biomedical Investigations(39). To balance completeness with usability, a practical path forward would be a tiered metadata model, in which a minimal set of required fields ensures basic interoperability, while an extended optional tier captures richer biological and experimental context when available. Defining these tiers, and agreeing on which sources of variation are essential versus optional, remains an important community standardization challenge.

### Future consortium expansion and directions

The scFAIR consortium aims to expand both the number of resources connected to the portal and the number of partner organizations participating in the definition, implementation, and evaluation of shared standards. Future integrations will improve cross resource query capabilities through sc fair.org, while also allowing additional repositories and knowledgebases to contribute to the scFAIR forum and help shape community recommendations for metadata, validation, and reproducibility.

Given the current composition of the consortium, scFAIR remains primarily focused on metazoan single cell data. However, the schema developments described here, including broader support for Ensembl resources and explicit reporting of genome assembly and annotation versions, provide a foundation for future expansion beyond animals. In particular, collaboration with plant, fungal, protist, or microbial single cell communities could help define which parts of the current schema are broadly reusable and which metadata elements require domain specific extensions, such as organism specific anatomy, developmental stage, strain, cultivar, isolate, growth condition, or environmental context.

A major future role for scFAIR could also be the coordination of curation efforts across resources. Manual curation of cell types, anatomical structures, developmental stages, and annotation evidence remains a major bottleneck in making single cell datasets FAIR. Analogous to the way the GO consortium coordinates gene product annotation across participating resources(40), scFAIR could provide a forum for distributing curation effort, harmonizing annotation practices, sharing ontology mappings, and avoiding duplicated work across single cell platforms. Such coordination would be particularly valuable for large atlases and cross species studies, where consistent ontology use and transparent annotation evidence are essential for reliable data reuse.

## Data and materials availability statement

scFAIR metadata schema is kept updated on the scFAIR schema github repository and listed on the https://www.sc-fair.org/metadata-schema page. All source and code used for this manuscript, including figures and tables scripts are deposited on this github repository. The scFAIR website is accessible through this URL: https://www.sc-fair.org/, and its source code is openly accessible at this github repo: scFAIR portal github repository.

## Acknowledgements

We would like to thank Luka Domitrovic for the web development of the sc-fair.org web portal. We also would like to thank Rachel Marcone and Ismail Ugur Bayindir for their inputs.

